# Controlling GRF4-GIF1 Expression for Efficient, Genotype-Independent Transformation Across Wheat Cultivars

**DOI:** 10.1101/2025.08.29.673042

**Authors:** Sadiye Hayta, Mark A Smedley, Meltem Bayraktar, Macarena Forner, Anna Backhaus, Clare Lister, Martha Clarke, Cristobal Uauy, Simon Griffiths

## Abstract

Bread wheat (*Triticum aestivum* L.) plays a vital role in global food security, and its continuous genetic improvement is essential to meet the demands of a rapidly growing world population. Advances in genome sequencing and assembly have positioned wheat as a model crop for functional genomics and have increased the demand for highly efficient, genotype-independent transformation systems. A fusion technology involving a GROWTH-REGULATING FACTOR (GRF) and a GRF-INTERACTING FACTOR (GIF) has emerged as a powerful tool to enhance regeneration efficiency and expand the range of transformable genotypes. In this study, we present an optimized and robust *Agrobacterium*-mediated wheat transformation protocol incorporating *GRF4-GIF1*, tested across multiple wheat varieties. Across all tested wheat cultivars, *GRF4-GIF1* containing constructs consistently enabled successful transformation, with varied efficiencies depending on the genotype and promotors used to drive the gene fusion. Our method significantly improves transformation efficiency while minimizing *GRF4-GIF1* pleiotropic effects, providing a versatile platform for gene function analysis and gene editing. This work represents a critical step toward efficient, genotype-independent transformation in wheat, supporting both research and breeding applications aimed at improving crop resilience, nutritional value, and productivity.

## Introduction

Bread wheat (*Triticum aestivum* L.), also known as common wheat, is one of the “big three” cereal crops worldwide (Shewry, 2009). With its extensive cultivation range and vital role as the primary source of cereal-based processed products. Wheat significantly contributes to global food security, providing approximately 20% of the world’s caloric intake and 25% of daily protein consumption. Furthermore, wheat is an important source of essential minerals for human nutrition and contributes up to 20% of essential dietary minerals in the UK (Sigalas *et al*., 2024). Improving wheat genetics to enhance its nutritional value, disease resistance, and resilience to climate change will have a profound impact on sustainable food production and global nutritional security.

The advent of advanced technologies, including the development of multiple reference-quality genome assemblies has ushered in a new era for bread wheat research, equipping researchers and breeders with vital tools to enhance wheat and address future food security challenges (Walkowiak *et al*., 2020). These genome assemblies provide a foundation for functional gene discovery and breeding, facilitating the development of the next generation of modern wheat cultivars. The availability of these genomic sequences, coupled with new open-access transformation protocols (Hayta *et al*., 2019), will enable the powerful application of molecular tools in future wheat research and breeding. Effective transgenic methodologies are now crucial for conducting gene functional studies, enhancing traits, and integrating precision breeding techniques.

Wheat transformation is highly genotype-dependent and typically relies on a few cultivars, such as Fielder and Bobwhite, which exhibit a strong tissue culture response (Hayta *et al*., 2019). Despite the first successful reports of *Agrobacterium*-mediated wheat transformation in the 1990s (Cheng *et al*., 1997), transformation efficiencies remained low—around 5%—for many years. An in-planta method was reported in 2009 (Risacher *et al*., 2009), but it was not widely adopted. Later, Japan Tobacco Inc. demonstrated improved transformation efficiency in the model wheat genotype Fielder using a patented, licensed system called PureWheat, which requires specialist training and specific vectors, thereby limiting its accessibility (Ishida *et al*., 2015).

Our first wheat CRISPR study, on the ZIP4-B2 (Ph1 locus) a meiotic gene involved in crossover within wheat (Rey *et al*., 2018), lead to the publication of our freely available, efficient, and reproducible transformation protocol in 2019, achieving transformation efficiencies averaging around 25% in Fielder, and also Kronos and Cadenza with lower efficiencies 10% and 4% respectively (Hayta *et al*., 2021). Kronos and Cadenza are important reference varieties, with extensive TILLING resources available for gene discovery and complementation (Krasileva *et al*., 2017). The key factors influencing wheat transformation include the cultivar used, the quality and health of the donor material, the developmental stage of the immature embryos, the handling of the material, and the media component composition, transformation efficiencies are drastically affected by multiple factors with narrow optimal windows (Hayta *et al*., 2021).

The genotype-independence of transformation techniques further broadens the potential for genetic improvements, making wheat breeding more versatile and accessible. Debernardi *et al*. (2020) developed a growth regulator fusion technology involving GROWTH-REGULATING FACTOR 4 (GRF4) and its cofactor GRF-INTERACTING FACTOR 1 (GIF1). Overexpressing *GRF4-GIF1* in wheat significantly boosts regeneration efficiency and broadens the range of transformable wheat genotypes. We tested this technology using our published wheat transformation method. Fielder plants transformed with the *GRF4–GIF1* chimera increased dramatically to 77.5% efficiency (Debernardi *et al*., 2020). This technology accelerates the transformation pipeline and enables high transformation efficiencies by overcoming narrow optimal windows, such as specific growing conditions and embryo developmental stages, which are critical for successful transformation in wheat.

Low transgene copy number *GRF4-GIF1* transgenic plants exhibited normal development and fertility in Fielder, however, constitutive expression of other plant developmental regulators such as the maize *Baby boom* (*ZmBbm*), the maize *Wuschel2* (*ZmWus2*) (Johnson *et al*., 2023), and a Wuschel homolog TaWox5 (Wang *et al*., 2022) can lead to negative pleiotropic effects, necessitating their removal from transgenic plants and limiting their practical application (Wang *et al*., 2020). In an effort to minimize the negative pleiotropic effects associated with developmental regulators, Lowe *et al*. (2016) demonstrated a strategy for successful plant regeneration using Cre recombinase, which excised morphogenic gene sequences flanked by loxP sites. This approach was critical in maize transformation, removing unwanted expression cassettes (*Bbm* and *Wus2*) linked to undesirable phenotypes, such as thick, short roots, and ensuring proper plant regeneration. Further advancements were made by Lowe *et al*. (2018) who identified the *ZmPLTP* (maize phospholipid transferase protein gene), a maize promoter with strong expression in leaves, embryos, and callus, but downregulated in roots, meristems, and reproductive tissues. This promoter was used to drive Bbm expression in maize, enabling efficient somatic embryo formation without the callus phase and rapid plant regeneration. The *ZmPLTP* allowed for uniform transformation and successful germination of maize plantlets. Co-transformation has been widely used because it is a simple and clean technique, leaving no residual DNA sequences, such as inverted repeats and recombination sites, in transgenic plants from which the selectable marker gene has been eliminated with high frequency. Co-transformation involves the simultaneous integration of a selectable marker gene and a gene of interest from different T-DNAs, followed by their subsequent recombination and segregation in the progeny, provided the two genes are integrated into unlinked loci.

Debernardi *et al*. (2020) demonstrated that transgenic plants overexpressing the *GRF4-GIF1* chimera under the control of the *ZmUbi* promoter maintained normal fertility and morphology. However, these plants exhibited a 23.9% reduction in grain number per spike and a 13.7% increase in grain weight, suggesting that *GRF4-GIF1* can modulate physiological traits without drastically impairing plant development. In our study, high transgene copy numbers of *GRF4-GIF1* were associated with varying degrees of sterility, depending on the cultivar, but also showed a modest positive effect on grain area and grain length. For effective trait gene function characterization, it is critical to develop transgenic plants that exhibit desirable traits while minimizing adverse effects linked to morphological gene overexpression. The *GRF4–GIF1* technology addresses genotype dependence and expands the applicability of transformation methods, including gene-editing tools, across elite wheat cultivars. Moreover, the ability to segregate or remove *GRF4–GIF1* after transformation provides enhanced flexibility for downstream applications such as functional genomics, gene editing and precision breeding. Importantly, *GRF4–GIF1* technology, together with open protocols, further democratises gene editing by making these approaches increasingly accessible to research centres and wheat breeders lacking high-end facilities.

In this study, we harness *GRF4-GIF1* technology to develop an optimized and highly efficient transformation protocol, validated across multiple wheat cultivars. Additionally, we propose strategies to mitigate undesirable pleiotropic effect such as, reduced grain number, thereby enhancing the accuracy and reliability of functional analyses for trait-associated genes.

## Results

### GRF Transcription Factors and Physiological Impact

GRF transcription factors are highly conserved across angiosperms, gymnosperms, and mosses, encoding proteins with conserved QLQ and WRC domains essential for protein-protein and protein-DNA interactions. In many plant lineages, GRFs are regulated by microRNA miR396, which downregulates GRF expression in mature tissues.

We first tested the *ZmUbi GRF4-GIF1* construct across nine hexaploid wheat cultivars (Table 1), including several that had previously been considered recalcitrant to transformation. A GUS-only construct served as the control. For *GRF4-GIF1* transformations, 70–100 immature embryos were used per cultivar, and 50–80 embryos were used for controls. Calli transformed with *ZmUbi*::*GRF4-GIF1* exhibited strong regenerative potential, demonstrating the construct’s ability to promote somatic embryogenesis (Figure 1).

**Table 1.**
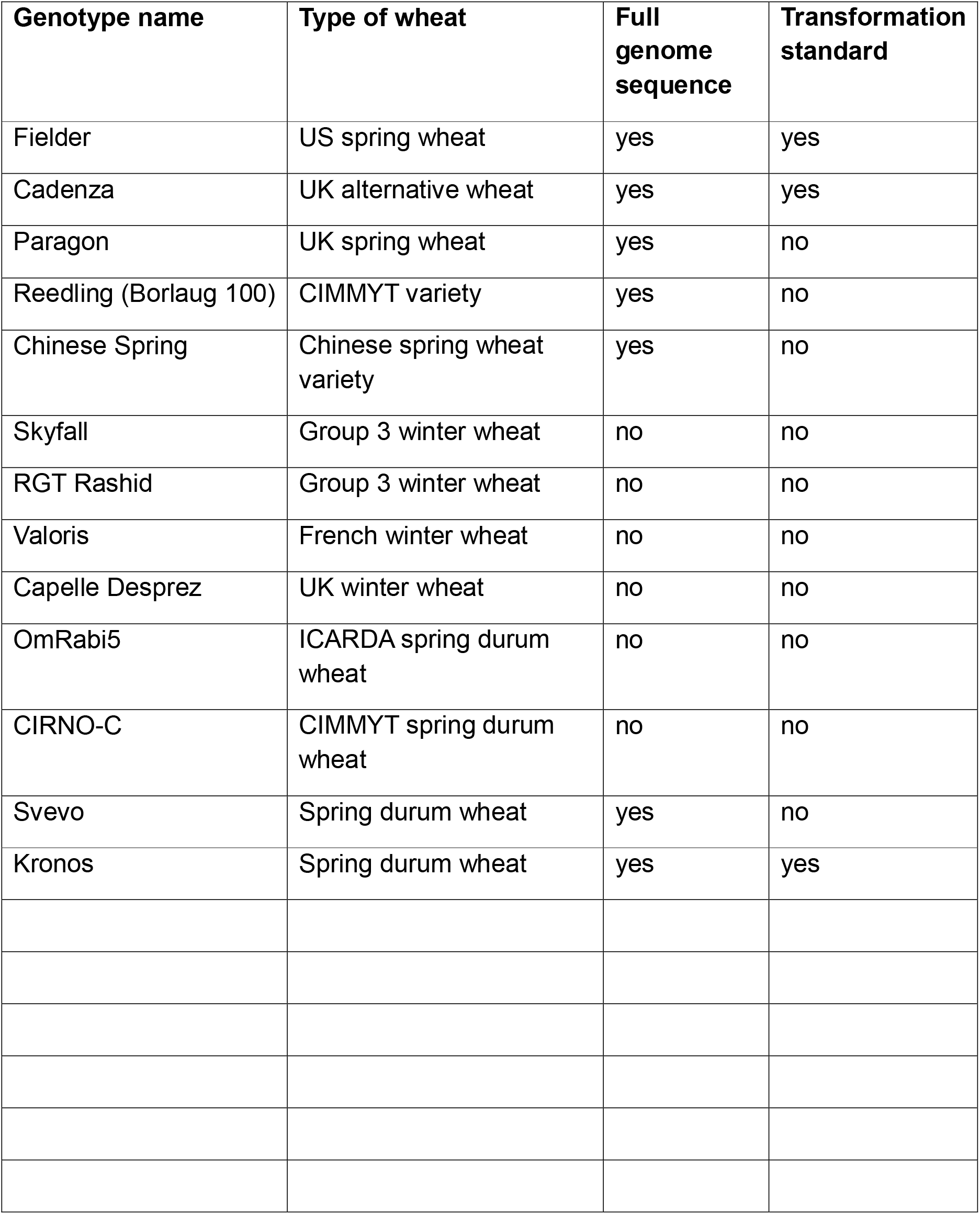
Hexaploid and tetraploid wheat cultivars used in transformation experiments. The table includes each cultivar’s classification, genome sequence availability, and whether it has been used as a transformation standard.

**Figure 1.**
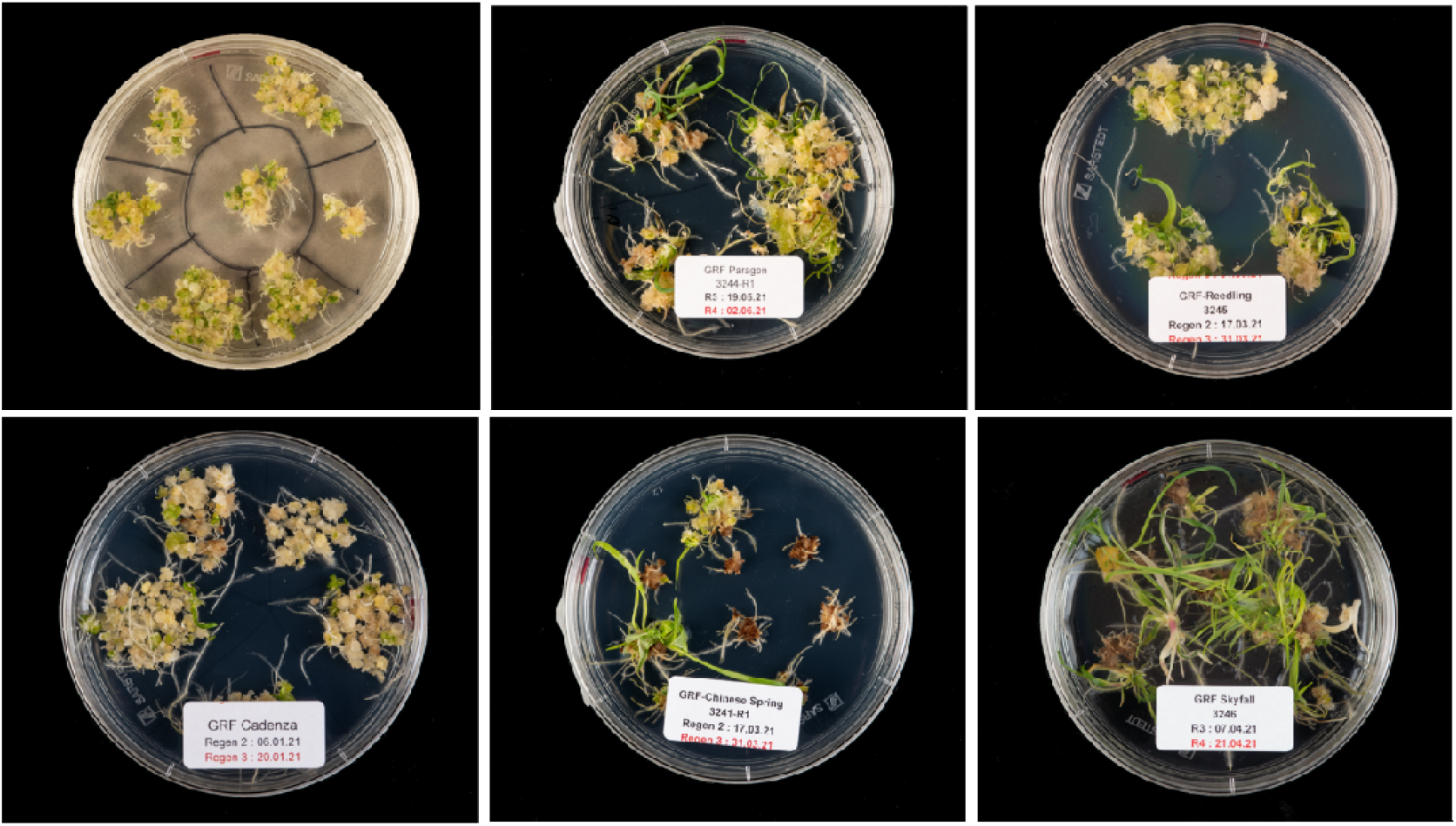
Regeneration of wheat calli using the *ZmUbi GRF4-GIF1* construct across different cultivars. Representative regeneration responses are shown for the wheat cultivars Fielder, Reedling (Borlaug 100), Cadenza, Chinese Spring, and Skyfall, respectively.

Our data from mature plants showed that the number of fertile spikelets was reduced by *GRF4-GIF1* expression across all varieties. Additionally, *GRF4-GIF1* lines produced fewer spikelets per plant than the controls. Lines with a high transgene copy number showed a slightly higher proportion of sterility, the severity of which depended on the cultivar. Among the cultivars tested, Kronos had the lowest seed set, indicating the greatest sensitivity to *GRF4-GIF1* whereas Paragon and Cadenza were less affected. *GRF4-GIF1* had a slightly positive effect on grain area and grain length across all cultivars, consistent with findings by Debernardi *et al*. (2020). A significant increase in the number of spikes was observed in Fielder, highlighting a strong tillering response in this background.

Variation in seed number among Paragon plants with differing *GRF4-GIF1* copy numbers (Fig. 2) underscores the impact of *GRF4-GIF1* overexpression on seed set. In gene editing studies, segregation of the T-DNA allows pleiotropic effects to be avoided in subsequent generations, enabling recovery of edited lines; however, in other transformation studies, particularly those involving sensitive traits, careful management is required.

**Figure 2.**
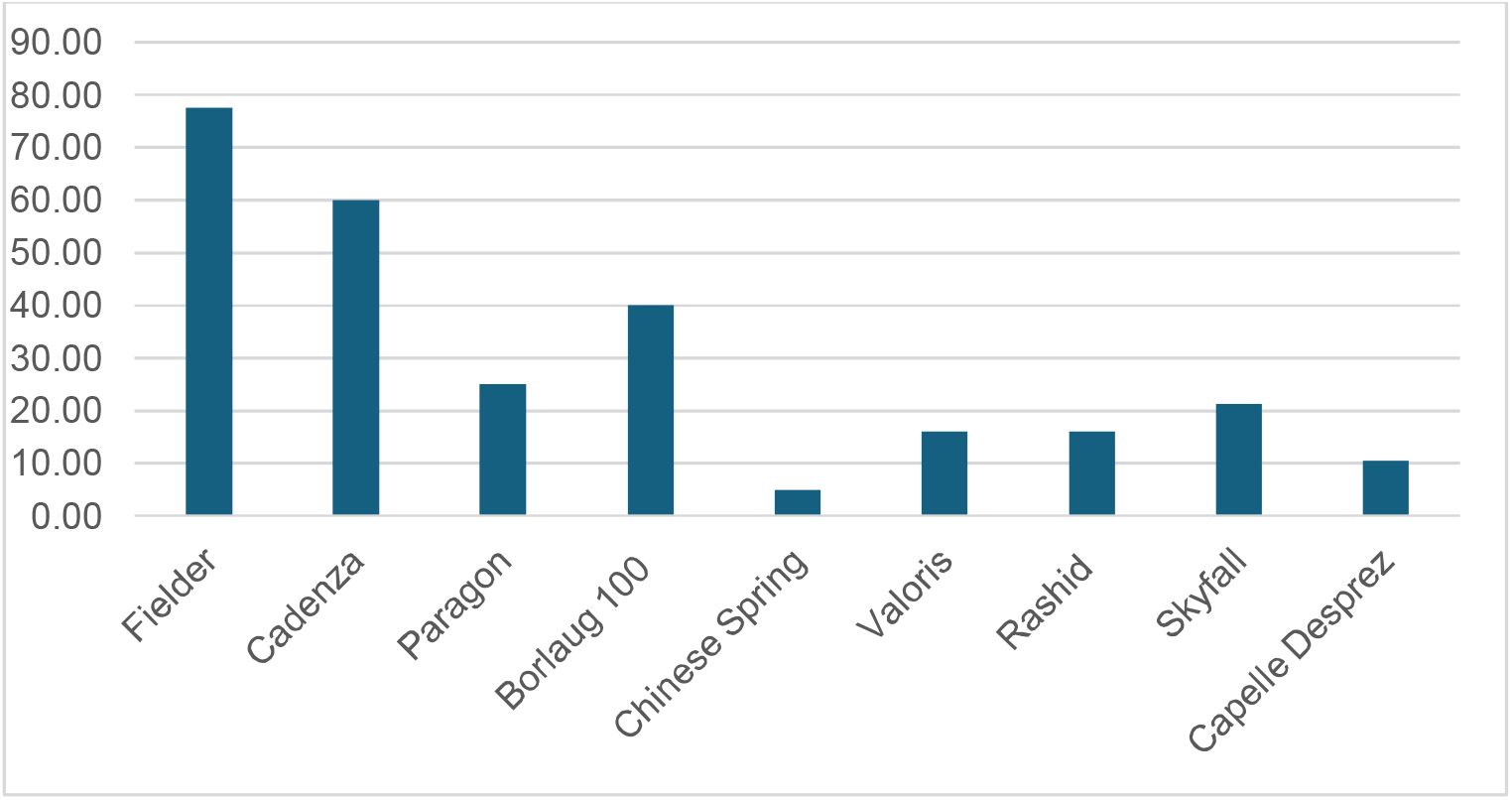
Transformation efficiencies in various wheat cultivars. Transformation efficiencies achieved using different GRF4-GIF1 constructs across multiple wheat cultivars, including elite and commercial lines.

### Transformation Efficiency and Phenotypic Effects in Hexaploid Wheat

Regeneration capacity with the *ZmUbi*-driven *GRF4-GIF1* construct was high, highlighting the effectiveness of *GRF4-GIF1* in promoting somatic embryogenesis in wheat (Fig. 1). Transformation efficiencies reached 77.5% in Fielder, 60% in Cadenza, and 40% in Borlaug 100. In winter wheat cultivars such as Valoris and Skyfall, efficiencies were 16% and 20%, respectively (Fig. 3). Notably, multiple transgenic plants were frequently regenerated from a single embryo, and approximately 80% of these transgenics displayed different transgene copy numbers, suggesting that the actual transformation efficiency may be even higher than reported. Only Fielder and Cadenza were amenable to transformation without the *GRF4-GIF1* fusion, albeit with substantially lower efficiencies, 25% and 5% respectively.

**Figure 3.**
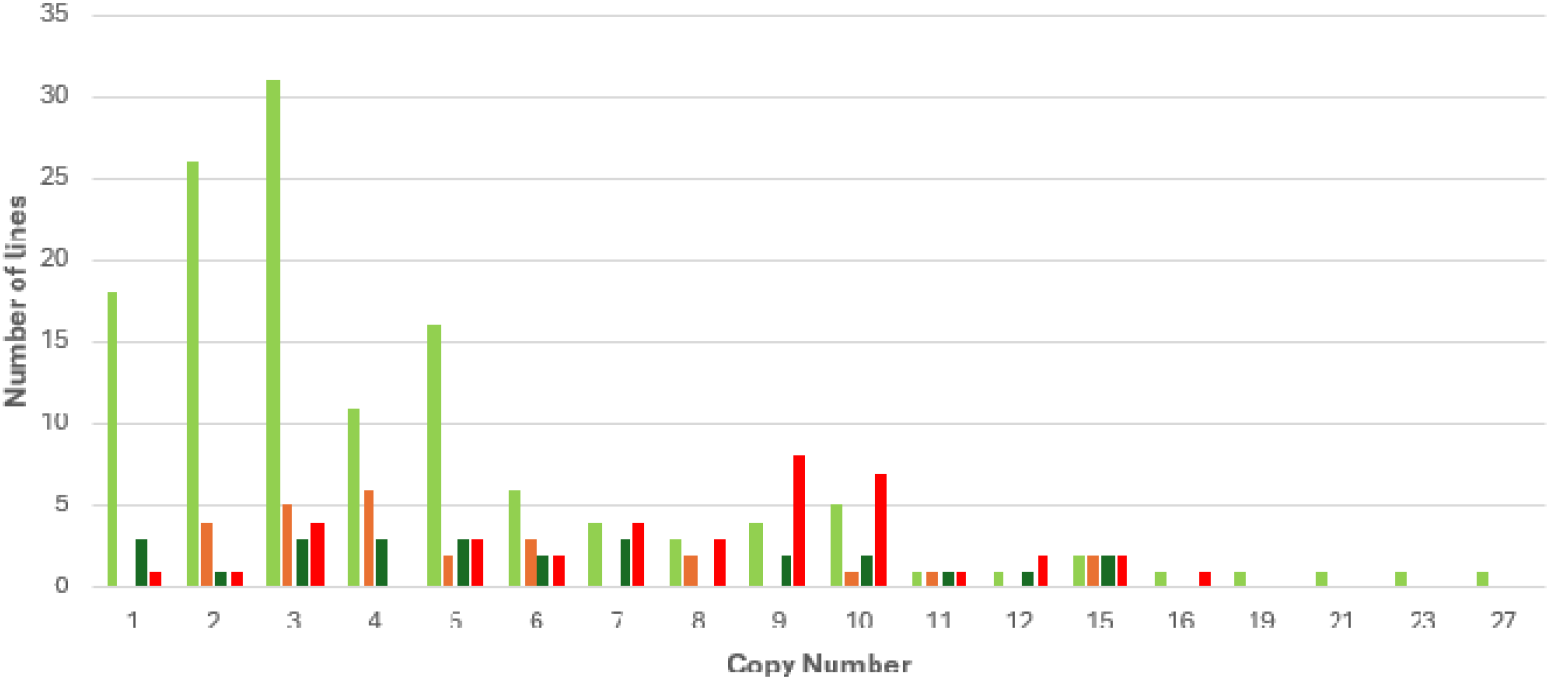
Variation in seed number among Paragon plants with different GRF4-GIF1 copy numbers. Bar colours indicate seed set phenotype categories: **Light green**: Number of plants with normal or near-normal seed number **Orange**: Number of plants with moderately reduced seed number **Dark green:** Number of plants with severely reduced seed number **Red**: Number of sterile plants

**Figure 4.**
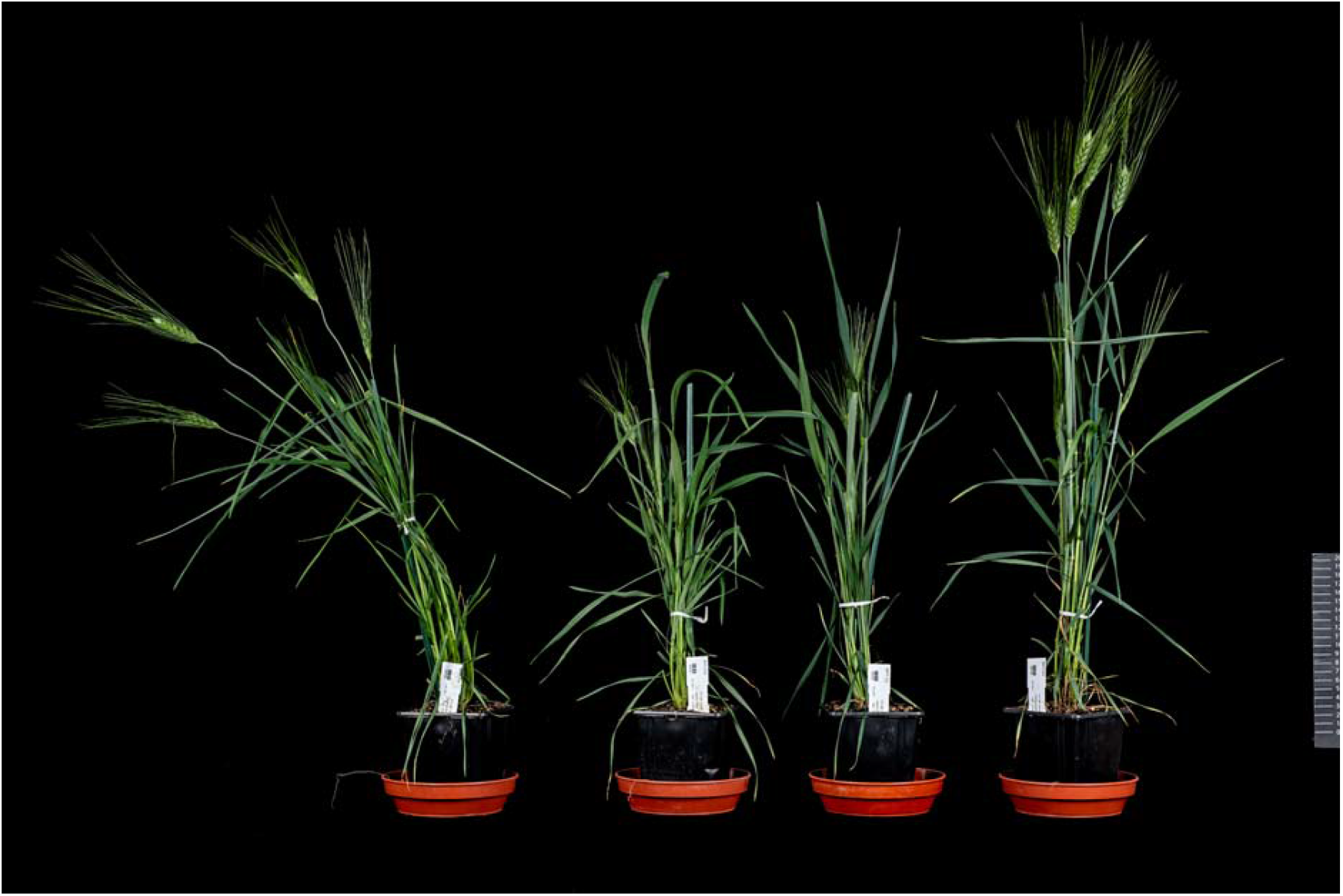
Phenotype of Kronos PLTP GRF4-GIF1 and non-transformed control plants (right). Transgenic Kronos plants expressing the ZmPLTP GRF4-GIF1 construct showed phenotypes closely resembling non-transformed controls, indicating minimal pleiotropic effects from transgene expression.

This advancement has enabled successful transformation across a broader range of wheat genotypes, including elite and commercial cultivars such as Borlaug 100, RGT Rashid, Skyfall, and Valoris. Among these, Skyfall is the most widely cultivated hexaploid bread making wheat in the UK, Valoris is a French variety notable for its high pentosan content, and Borlaug 100, developed by CIMMYT, serves as a key international breeding line. By overcoming the narrow optimal transformation windows typically associated with these genotypes, the *GRF4-GIF1* technology has achieved high transformation efficiencies while preserving protocol flexibility. Importantly, the method accommodates embryos of varying sizes sourced from plants grown under different environmental conditions, thereby reducing the overall transformation timeline. Across all tested wheat cultivars, *GRF4-GIF1* constructs consistently enabled successful transformation, although efficiency varied depending on the genotype and construct used. Skyfall, RGT Rashid, Valoris, and Cappelle Desprez are winter wheat varieties, and as such, are traditionally considered more recalcitrant to transformation than Spring varieties, these varieties, all showed lower transformation efficiencies.

### Extending *GRF4-GIF1* to Tetraploid Wheat

Based on our initial observations indicating an increased sensitivity of tetraploid wheat cultivar, Kronos to *GRF4-GIF1* overexpression, we developed a series of constructs designed to either regulate its expression or facilitate its excision (Table 2).

**Table 2:** Constructs Used in Different Tetraploid Wheat Cultivars.

| Construct name | Promoter driving GRF4-GIF1 | Reason for choosing | Marker Gene | Addgene Number |
| --- | --- | --- | --- | --- |
| pGGG ZmUbi GRF4-GIF1 | ZmUbi:: *TaGRF4-GIF1* | Strong promoter | OsUbi:: *GUS* | # 246478 |
| pGGG OsAct GRF4-GIF1 | OsAct:: *TaGRF4-GIF1* | Medium promoter | OsUbi:: *GUS* | # 246479 |
| pGGG 35S GRF4-GIF1 | CaMV35S:: *TaGRF4-GIF1* | Weak promoter | OsUbi:: *GUS* | # 246480 |
| pGGG HSCre GRF4-GIF1 | ZmUbi:: *TaGRF4-GIF1* | Excisable via Heat Shock | OsUbi:: *GUS* | # 246481 |
| pGGG PLTP GRF4-GIF1 | ZmPLTP:: *TaGRF4-GIF1* | Tissue-specific promoter | OsUbi:: *GUS* | # 246482 |
| pGGG GRF5-GIF1 | ZmUbi:: *TaGRF5-GIF1* | Strong promoter | OsUbi:: *GUS* | # 246483 |
| pGGG GUS | Non-GRF4-GIF1 Control | Control | OsUbi:: *GUS* | # 165418 |
| pSoup HSCre GRF4-GIF1 | ZmUbi:: *TaGRF4-GIF1* | Excisable via Heat Shock | N/A | # 246484 |

These constructs enabled us to explore strategies for minimizing phenotypic abnormalities while maintaining transformation efficiency, thus tailoring the system to the specific needs of tetraploid wheat. Expanding our analysis to tetraploid wheat cultivars, in addition to Kronos, (TILLING mutant population available), we tested the constructs in several other cultivars, including CIRNO-C (developed by CIMMYT), Omrabi5 (from ICARDA), and Svevo (a well-known Italian durum wheat cultivar).

Transformation efficiencies varied significantly across the tetraploid wheat cultivars tested. The Italian durum wheat variety Svevo exhibited the highest efficiency (55%) using the *ZmUbi GRF4-GIF1* construct, while Kronos, CIRNO-C and Om Rabi showed lower efficiencies at 50%, 11% and 4%, respectively. As expected, transformation outcomes varied depending on both the construct used and the cultivar. Notably, the highest efficiencies across all tetraploid lines were consistently achieved when the *GRF4-GIF1* fusion was driven by the strong *ZmUbi* promoter.

In many plant transformation systems, cytokinins are essential for shoot regeneration, with zeatin being the preferred cytokinin in wheat transformation. Previous high-efficiency protocols have typically used zeatin at 5 mg/L (Ishida, 2015). In our study, Svevo, which exhibited excellent regeneration capacity, achieved a transformation efficiency of 55% using the *GRF4-GIF1* construct with only 1.0 mg/L zeatin, indicating that the *GRF4-GIF1* chimera can promote embryogenesis, shoot proliferation, or both without the need for high zeatin concentrations. This finding aligns with Debernardi *et al*. (2020), who also reported high transformation efficiencies in Fielder without the addition of exogenous cytokinin. Svevo maintained a moderate transformation efficiency of 15% when transformed with the control construct lacking *GRF4-GIF1*, while Kronos showed 8.5% efficiency under the same conditions. In contrast, cultivars such as CIRNO-C and Kronos demonstrated improved transformation outcomes when cultured with a higher zeatin concentration of 2.5 mg/L, with Kronos reaching 50% efficiency. Notably, CIRNO-C could not be transformed without the *GRF4-GIF1* construct. All cultivars produced more plantlets per immature embryo when cultured with 2.5 mg/L zeatin. These results highlight the genotype-specific hormonal requirements and underscore the critical role of *GRF4-GIF1* in enabling the transformation of otherwise recalcitrant tetraploid wheat lines.

To further mitigate phenotypic effects, particularly sterility associated with high transgene copy numbers, we employed two complementary strategies: (1) using the non-constitutive *ZmPLTP* promoter to drive tissue-specific expression of *GRF4-GIF1*, restricting its activity to embryogenic tissues during early transformation stages; and (2) implementing a Cre/lox-mediated excision system with heat-inducible Cre recombinase to remove the *GRF4-GIF1* cassette. These approaches were evaluated in both the hexaploid cultivar Fielder and the tetraploid cultivar Kronos.

### Tissue-Specific Expression of *GRF4-GIF1* to Minimize Phenotypic Effects

In Fielder, the *ZmUbi GRF4-GIF1* construct delivered the highest transformation efficiency at 62%, producing an average of 3.1 plantlets per embryo. We measured tiller numbers, booting, and heading time after acclimatization. However, this construct also induced pronounced pleiotropic effects, including a high tiller count of 26.1 at week 5, compared to 11.8 in non-transformed controls. This increase in tiller number was further confirmed by comprehensive data collected from mature plants. By contrast, the *ZmPLTP GRF4-GIF1* construct achieved a transformation efficiency of 20%, generating 1.7 plantlets per embryo with a low escape rate of 0.4%. More importantly, these plants produced only 7.8 tillers at maturity, a substantial reduction of an undesirable pleiotropic effect.

In Kronos, the *ZmPLTP GRF4-GIF1* construct yielded a 23% transformation efficiency and an 11% escape rate, with a marked improvement in seed number and normal plant development. On average, transgenic plants produced 6.3 tillers and initiated flowering within five weeks, closely resembling the developmental timeline of non-transformed controls. Kronos benefited more significantly from the *ZmPLTP GRF4-GIF1* construct, demonstrating its suitability for minimizing phenotypic disruptions in sensitive genotypes.

Together, these results underscore the importance of promoter selection in wheat transformation. Tissue-specific expression driven by the *ZmPLTP* promoter enables efficient transformation while minimizing undesirable phenotypic effects (Figure 3).

### Excision of GRF4-GIF1 via Heat-Induced Cre-Lox System

In Kronos, successful *GRF4-GIF1* excision was observed in 11 out of 15 plants (73%), while in Fielder, excision occurred in 8 out of 20 plants (40%). The phenotype of excised plants for traits such as reduced tillering and improved fertility was closer to that of untransformed controls while those with the *ZmUbi*-driven *GRF4-GIF1* construct continued to display stronger developmental deviations. These outcomes are consistent with excision-based improvements previously reported in maize (Wang *et al*., 2020), which led to reduced ZmBbm and ZmWus2 morphological genes and fertile T0 plants (Aesaert *et al*., 2022) further supporting the utility of this system in mitigating *GRF4-GIF1* phenotypic associated effects.

### Co-Transformation and Comparison with Other Morphogenic Genes

We explored co-transformation as a method to introduce both *GRF4-GIF1* and other genes of interest (GOIs) into wheat. By using a two-*Agrobacterium* system with separate T-DNAs for *GRF4-GIF1* and the GOI, we achieved transformation efficiencies comparable to those obtained with the *ZmUbi GRF4-GIF1* construct. This approach offers greater flexibility and the potential for segregation of *GRF4-GIF1* and GOI in subsequent generations. We applied this strategy in elite wheat cultivars; Borlaug 100, Skyfall, Kronos, and Cadenza, demonstrating that constructs with two separate T-DNAs effectively introduced existing transgenes, achieving transformation efficiencies comparable to those obtained with the *ZmUbi GRF4-GIF1* construct.

We also compared *GRF4-GIF1* with other morphogenic genes, including WOX5. Wang and colleagues (2022) reported that the wheat homolog of Arabidopsis WUSCHEL-RELATED HOMEOBOX 5 (WOX5) enhances wheat transformation efficiency and promotes genotype independence. We tested the *ZmUbi*::WOX5 construct in both Fielder and Kronos cultivars. In Fielder, transformation with *ZmUbi*::WOX5 resulted in a 20% independent event rate and produced an average of 1.04 plantlets per line. In Kronos, the independent event rate was slightly higher at 24%. However, mature WOX5-transformed plants displayed notable phenotypic changes, including shorter stature, larger grain size, and wider, spiralled leaves (Fig. 5). In contrast, *GRF4-GIF1* consistently achieved higher transformation efficiency with fewer undesirable phenotypic alterations.

**Figure 5.**
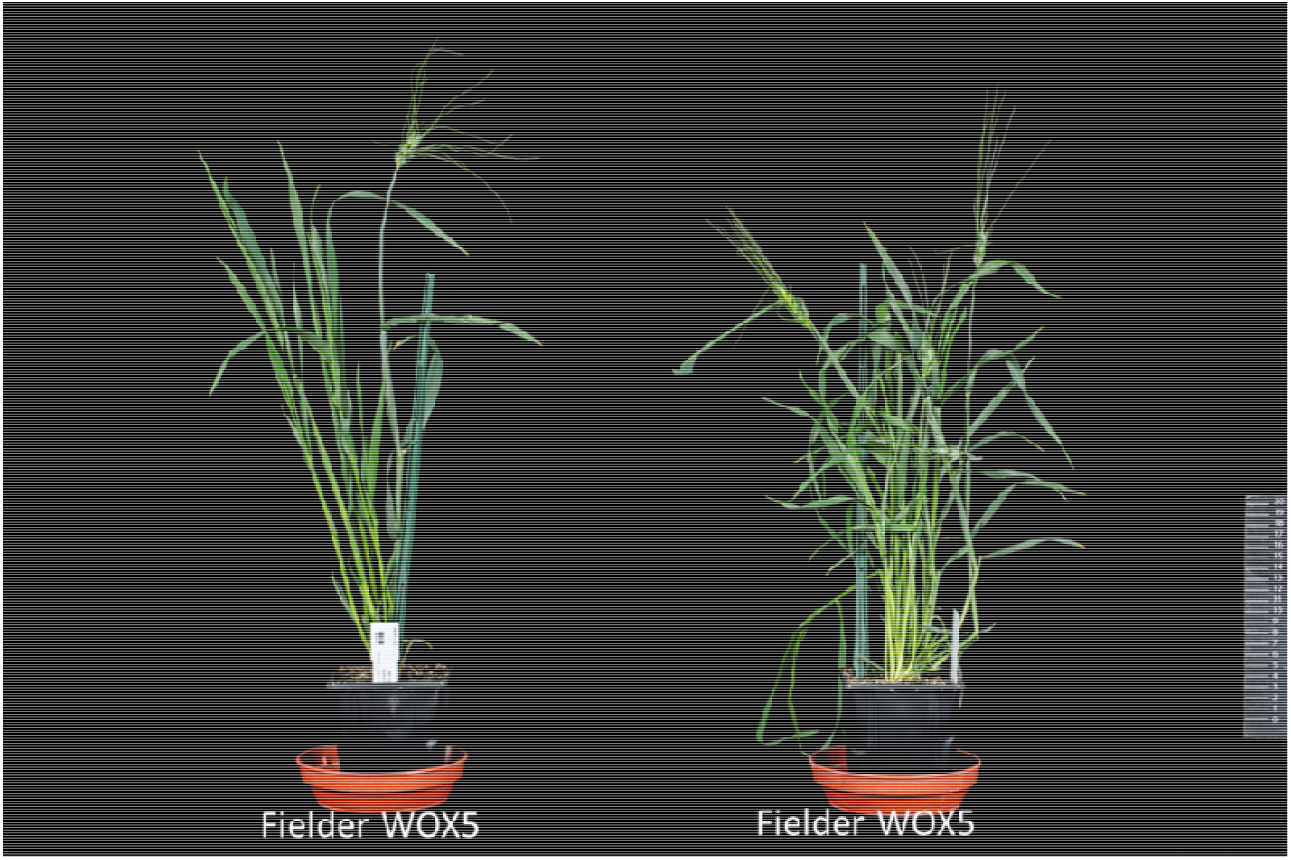
Phenotype of Fielder plants transformed with Wox5. Fielder plants expressing Wox5 exhibited distinct morphological changes compared to non-transformed controls. The mature plants displayed a shorter stature, larger grain size, and wider, spiralled leaf shapes, indicating significant developmental effects associated with Wox5 overexpression.

## Conclusion

Our study evaluated the effects of *GRF4-GIF1* constructs, driven by promoters with varying strength and temporal activity, in combination with different growth regulators across multiple tetraploid and hexaploid wheat cultivars. By assessing both transformation efficiency and developmental outcomes, we demonstrated that incorporation of *GRF4-GIF1* into wheat transformation protocols substantially improves efficiency and broadens the spectrum of amenable genotypes. Furthermore, the strategic use of promoters such as *ZmPLTP* and post-transformation excision systems like heat-inducible Cre/lox minimised undesirable phenotypic effects while preserving transformation robustness. Together, these advances provide an effective framework for precise gene editing in elite cultivars, facilitating streamlined integration into precision breeding pipelines.

## Material

### Plant Material and Growth Conditions

In this study, we utilized selected hexaploid bread wheat (*Triticum aestivum* L.) cultivars, as well as a few tetraploid durum wheat (Triticum durum Desf.) cultivars (Table 1). The different wheat cultivars were successive sown and grown under the growing conditions described in Hayta *et al*., 2021. Winter wheat cultivars were exposed to a 4-week vernalisation period at temperature 5°C, under a short-day, 8 h light:16 h dark photoperiod, 70 µmol m^-2^ s^-1^.

### Plasmid Constructs

The hypervirulent *Agrobacterium* tumefaciens strain AGL1 (Lazo *et al*., 1991) was used in all plant transformation experiments. All transformation vectors were based on the binary vector pGoldenGreenGate (pGGG) (Smedley *et al*., 2021) and were assembled using MoClo golden gate assembly (Weber *et al*., 2011). Each pGGG vector contained the hygromycin selectable marker (hpt) and Cat1 intron driven by the switchgrass ubiquitin promoter (Addgene # 246468), the *GRF4-GIF1* gene fusion driven by various promoters or excisable and the β-glucuronidase gene (GUS) with two introns (GUS2Int) under the control of the rice ubiquitin promoter (Addgene # 246471). The pGGG vectors were electroporated into the *Agrobacterium* AGL1 competent cells with the helper plasmid pAL155 (Addgene # 246472) which contains an additional VirG gene as previously described in Hayta *et al*. (2019).

The *GRF4-GIF1* gene fusion was synthesised as a MoClo golden gate level 0 component, internal Eco31I and BpiI restriction enzyme sites were removed by silent mutation and standardised MoClo overhangs and Eco31I sites added for assembly. The level 0 *GRF4-GIF1* CDS was assembled into expression cassettes in the Level 1 Position 2 pICH47742 (Addgene # 48001) with the promoters listed in Table 2 and the Nos terminator. All the regular promoters listed were at our laboratory’s disposal, however, the Zea mays phospholipid-transfer protein (*ZmPLTP*) promoter was designed and synthesised as a MoClo golden gate Level 0 component, internal Eco31I and BpiI restriction enzyme sites removed and standardised MoClo overhangs and Eco31I sites added for assembly. The pGGG *GRF4-GIF1* GUS (Addgene # 246478) vector containing the maize ubiquitin promoter (*ZmUbi*) driving the *GRF4-GIF1* and the GUS marker gene served as a control.

The inducible site-specific recombinase (Cre) recombinase CDS under the control of the barley heat shock-inducible promoter 17 (HvHsp17) was deployed for conditional excision of the *ZmUbi*::*GRF4-GIF1*::NosT and self-excision of the HvHsp17::Cre::HSTerm. A loxP acceptor cassette was designed and synthesised (Genewiz), which enabled the insertion via BpiI of Level 1 expression cassettes flanked by loxP sites. The cassette in-turn could be isolated via Eco31L and cloned into any MoClo L1 acceptor plasmid. The L1P2 *HvHsp17*::*CreInt*::*HvHsp17*::Term and L1P1 *ZmUbi*::*GRF4-GIF1*::Nos were assembled into the loxP cassette, and then the whole assembly was inserted into pICH47811 (AddGene #48008), this vector was deemed L1P2 Rev Cre/LoxP heat shock excisable *GRF4-GIF1* (Addgene # 246486), before cloning in the reverse orientation upstream of the hpt selection cassette in the final Level 2 pGGG binary vector.

Constructs with two separate T-DNAs provide the opportunity to either use “old” constructs in elite wheat cultivars without having to re-cloning to insert *GRF4-GIF1* or/and to obtain *GRF4-GIF1*-free plants through segregation in later generations. A T-DNA cassette was designed and synthesised which was flanked by left and right border repeat sequences and was classically cloned into the pSoup helper plasmid (Addgene # 165419) via the KpnI and HindIII restriction enzyme sites. The L1P2 Cre loxP GRF-GIF Cassette (Addgene # 246486) was cloned into the pSoup’s T-DNA via Eco31L using a golden gate assembly. The resulting plasmid known as pSoup HSCre *GRF4-GIF1* (Addgene # 246484) can be co-electroporated into AGL1 with pGGG based plasmids or an *Agrobacterium* culture containing pSoup HSCre *GRF4-GIF1* can be mixed with an *Agrobacterium* culture containing a suitable plasmid of interest and co-transformed into wheat.

Various *GRF4-GIF1* constructs, driven by different promoters, were designed for testing in tetraploid wheat, which is more sensitive to phenotypic changes than hexaploid bread wheat. Additionally, *GRF5-GIF1* was included as an alternative construct to assess its potential in tetraploid wheat, alongside *TaWOX5*, which was tested in Kronos and Fielder.

### Agrobacterium-mediated Transformation of Different Wheat Cultivars

Embryos were isolated from the immature grains at the early milk stage GS73 and underwent *Agrobacterium*-mediated transformation following previously published protocols by (Hayta *et al*., 2019, Hayta *et al*., 2021) with modifications depending on the wheat cultivars. After transformation, calli were selected on media containing two levels of hygromycin, Selection 1 (S1) and Selection 2 (S2). They were subsequently transferred to the regeneration medium with defined concentrations of hygromycin and zeatin riboside, followed by culture on rooting media for root induction (Table 3).

**Table 3.** Selection and Regeneration Conditions for *in Vitro* Wheat Calli With or Without *GRF4-GIF1*.

| Wheat Cultivar | S1 Hygromycin (mg/L) | S2 Hygromycin (mg/L) | Regeneration Hygromycin (mg/L) | Zeatin (mg/L) |
| --- | --- | --- | --- | --- |
| Fielder | 20 | 30 | 20 | 2.5 |
| Fielder + GRF4-GIF1 | 15 | 15 | 15 | 1.0 |
| Kronos | 15 | 15 | 15 | 2.5 |
| Kronos + GRF4-GIF1 | 15 | 15 | 15 | 2.5 |
| Cadenza | 10 | 10 | 10 | 1.0 |
| Cadenza + GRF4-GIF1 | 10 | 10 | 10 | 1.0 |
| Skyfall + GRF4-GIF1 | 10 | 10 | 10 | 2.5 |
| Reedling + GRF4-GIF1 | 10 | 10 | 10 | 1.0 |
| Cappelle Desprez + GRF4-GIF1 | 10 | 10 | 10 | 1.0 |
| Paragon + GRF4-GIF1 | 5 | 5 | 5 | 1.0 |
| Valoris | 5 | 5 | 5 | 1.0 |
| Chinese Spring + GRF4-GIF1 | 5 | 5 | 5 | 2.5 |

The co-transformation system was tested in our transformation system. *Agrobacterium* containing our gene of interest (GOI) construct was co-transformed with *Agrobacterium* containing the pSoup HSCre *GRF4-GIF1* construct. The *Agrobacterium* overnight cultures were adjusted to an OD_600_ of 0.4 and then mixed at a 2:1 ratio of GOI to *GRF4-GIF1* prior to embryo inoculation. Alternatively, pGGG vectors lacking *GRF4-GIF1* were electroporated into *Agrobacterium* strains already containing pSoup HSCre *GRF4-GIF1*, and standard inocula were prepared and used for transformation as described by Hayta *et al*. (2019).

### Heat-Shock Treatment for Inducible Site-specific Recombinase (Cre) to Excise GRF4-GIF1

Plant material transformed with constructs containing the heat-shock-inducible Cre/lox-mediated excision system for *GRF4-GIF1* was subjected to a 40°C heat treatment for 16 hours (overnight) during the second regeneration phase. Only calli that had developed green shoots were exposed to heat-shock prior to subculturing onto fresh media for further development.

### DNA Extraction and Copy Number Analysis

Leaf samples 0.5 to 0.7 cm were collected, and DNA extracted using the protocol adapted from (Pallotta, 2003). Transgenesis was confirmed and transgene copy number analysis performed using Taqman qPCR and probe as described in Hayta *et al*., (2019). The values obtained were used to calculate transgene copy number using the 2^−ΔΔCT^ calculation as described in (Livak and Schmittgen, 2001).

The transformation efficiency was calculated as single events occurring from each embryogenic line (1 plant from 1 embryo) and displayed as the percentage of positive transgenic plants produced from the total number of immature embryos isolated and inoculated with *Agrobacterium* in an experiment.

### Data Collection

Full phenotypic traits, including plant height, tiller, spikelet numbers, grain per spike and thousands grain weight (TGW), were measured at maturation.

### Statistical Analysis

Data were analysed using ANOVA on phenotypic data in RStudio (v1.3.1056). Means were compared using Tukey’s HSD test with a significance level of p < 0.05.

## Acknowledgements

The authors acknowledge and thank Juan Debernardi for providing insightful advice. The authors also acknowledge and thank Mark Youles of TSL SynBio for supplying MoClo L0 components. This work was supported by the BBSRC Institute Strategic Programme (ISP) “Delivering Sustainable Wheat (DSW)” (BB/X011003/1), and the Scientific and Technological Research Council of Turkey (TUBITAK) through the 2219 – International Postdoctoral Research Fellowship Program.

## References

Aesaert, S., Impens, L., Coussens, G., Van Lerberge, E., Vanderhaeghen, R., Desmet, L., Vanhevel, Y., Bossuyt, S., Wambua, A.N., Van Lijsebettens, M., Inzé, D., De Keyser, E., Jacobs, T.B., Karimi, M. and Pauwels, L. (2022) Optimized Transformation and Gene Editing of the B104 Public Maize Inbred by Improved Tissue Culture and Use of Morphogenic Regulators. Front Plant Sci, Volume 13 - 2022.

Cheng, M., Fry, J.E., Pang, S., Zhou, H., Hironaka, C.M., Duncan, D.R., Conner, T.W. and Wan, Y. (1997) Genetic Transformation of Wheat Mediated by Agrobacterium tumefaciens. Plant Physiol, 115, 971–980.

Debernardi, J.M., Tricoli, D.M., Ercoli, M.F., Hayta, S., Ronald, P., Palatnik, J.F. and Dubcovsky, J. (2020) A GRF-GIF chimeric protein improves the regeneration efficiency of transgenic plants. Nat Biotechnol, 38, 1274–1279.

Hayta, S., Smedley, M.A., Clarke, M., Forner, M. and Harwood, W.A. (2021) An Efficient Agrobacterium-Mediated Transformation Protocol for Hexaploid and Tetraploid Wheat. Current Protocols, 1, e58.

Hayta, S., Smedley, M.A., Demir, S.U., Blundell, R., Hinchliffe, A., Atkinson, N. and Harwood, W.A. (2019) An efficient and reproducible Agrobacterium-mediated transformation method for hexaploid wheat (Triticum aestivum L.). Plant Methods, 15, 121.

Ishida, Y., Tsunashima, M., Hiei, Y. and Komari, T. (2015) Wheat (Triticum aestivum L.) transformation using immature embryos. Methods in molecular biology (Clifton, N.J.), 1223, 189–198.

Johnson, K., Cao Chu, U., Anthony, G., Wu, E., Che, P. and Jones, T.J. (2023) Rapid and highly efficient morphogenic gene-mediated hexaploid wheat transformation. Frontiers in Plant Science, 14.

Krasileva, K.V., Vasquez-Gross, H.A., Howell, T., Bailey, P., Paraiso, F., Clissold, L., Simmonds, J., Ramirez-Gonzalez, R.H., Wang, X., Borrill, P., Fosker, C., Ayling, S., Phillips, A.L., Uauy, C. and Dubcovsky, J. (2017) Uncovering hidden variation in polyploid wheat. Proc Natl Acad Sci U S A, 114, E913–e921.

Lazo, G.R., Stein, P.A. and Ludwig, R.A. (1991) A DNA transformation-competent Arabidopsis genomic library in Agrobacterium. Biotechnology (N Y), 9, 963–967.

Livak, K.J. and Schmittgen, T.D. (2001) Analysis of Relative Gene Expression Data Using Real-Time Quantitative PCR and the 2−ΔΔCT Method. Methods, 25, 402–408.

Lowe, K., La Rota, M., Hoerster, G., Hastings, C., Wang, N., Chamberlin, M., Wu, E., Jones, T. and Gordon-Kamm, W. (2018) Rapid genotype “independent” Zea mays L. (maize) transformation via direct somatic embryogenesis. In Vitro Cell Dev Biol Plant, 54, 240–252.

Lowe, K., Wu, E., Wang, N., Hoerster, G., Hastings, C., Cho, M.J., Scelonge, C., Lenderts, B., Chamberlin, M., Cushatt, J., Wang, L., Ryan, L., Khan, T., Chow-Yiu, J., Hua, W., Yu, M., Banh, J., Bao, Z., Brink, K., Igo, E., Rudrappa, B., Shamseer, P.M., Bruce, W., Newman, L., Shen, B., Zheng, P., Bidney, D., Falco, C., Register, J., Zhao, Z.Y., Xu, D., Jones, T. and Gordon-Kamm, W. (2016) Morphogenic Regulators Baby boom and Wuschel Improve Monocot Transformation. Plant Cell, 28, 1998–2015.

Pallotta, M., P Warner, RL Fox, H Kuchel, SJ Jefferies and P Langridge (2003) Marker assisted wheat breeding in the southern region of Australia. Proceedings of the Tenth International Wheat Genetics Symposium p.789–791.

Rey, M.D., Martín, A.C., Smedley, M., Hayta, S., Harwood, W., Shaw, P. and Moore, G. (2018) Magnesium Increases Homoeologous Crossover Frequency During Meiosis in ZIP4 (Ph1 Gene) Mutant Wheat-Wild Relative Hybrids. Front Plant Sci, 9, 509.

Risacher, T., Craze, M., Bowden, S., Paul, W. and Barsby, T. (2009) Highly efficient Agrobacterium-mediated transformation of wheat via in planta inoculation. Methods in molecular biology (Clifton, N.J.), 478, 115–124.

Shewry, P.R. (2009) Wheat. J Exp Bot, 60, 1537–1553.

Sigalas, P.P., Shewry, P.R., Riche, A., Wingen, L., Feng, C., Siluveru, A., Chayut, N., Burridge, A., Uauy, C., Castle, M., Parmar, S., Philp, C., Steele, D., Orford, S., Leverington-Waite, M., Cheng, S., Griffiths, S. and Hawkesford, M.J. (2024) Improving wheat grain composition for human health by constructing a QTL atlas for essential minerals. Communications Biology, 7, 1001.

Smedley, M.A., Hayta, S., Clarke, M. and Harwood, W.A. (2021) CRISPR-Cas9 Based Genome Editing in Wheat. Current Protocols, 1, e65.

Walkowiak, S., Gao, L., Monat, C., Haberer, G., Kassa, M.T., Brinton, J., Ramirez-Gonzalez, R.H., Kolodziej, M.C., Delorean, E., Thambugala, D., Klymiuk, V., Byrns, B., Gundlach, H., Bandi, V., Siri, J.N., Nilsen, K., Aquino, C., Himmelbach, A., Copetti, D., Ban, T., Venturini, L., Bevan, M., Clavijo, B., Koo, D.-H., Ens, J., Wiebe, K., N’Diaye, A., Fritz, A.K., Gutwin, C., Fiebig, A., Fosker, C., Fu, B.X., Accinelli, G.G., Gardner, K.A., Fradgley, N., Gutierrez-Gonzalez, J., Halstead-Nussloch, G., Hatakeyama, M., Koh, C.S., Deek, J., Costamagna, A.C., Fobert, P., Heavens, D., Kanamori, H., Kawaura, K., Kobayashi, F., Krasileva, K., Kuo, T., McKenzie, N., Murata, K., Nabeka, Y., Paape, T., Padmarasu, S., Percival-Alwyn, L., Kagale, S., Scholz, U., Sese, J., Juliana, P., Singh, R., Shimizu-Inatsugi, R., Swarbreck, D., Cockram, J., Budak, H., Tameshige, T., Tanaka, T., Tsuji, H., Wright, J., Wu, J., Steuernagel, B., Small, I., Cloutier, S., Keeble-Gagnère, G., Muehlbauer, G., Tibbets, J., Nasuda, S., Melonek, J., Hucl, P.J., Sharpe, A.G., Clark, M., Legg, E., Bharti, A., Langridge, P., Hall, A., Uauy, C., Mascher, M., Krattinger, S.G., Handa, H., Shimizu, K.K., Distelfeld, A., Chalmers, K., Keller, B., Mayer, K.F.X., Poland, J., Stein, N., McCartney, C.A., Spannagl, M., Wicker, T. and Pozniak, C.J. (2020) Multiple wheat genomes reveal global variation in modern breeding. Nature, 588, 277–283.

Wang, K., Shi, L., Liang, X., Zhao, P., Wang, W., Liu, J., Chang, Y., Hiei, Y., Yanagihara, C., Du, L., Ishida, Y. and Ye, X. (2022) The gene TaWOX5 overcomes genotype dependency in wheat genetic transformation. Nature Plants, 8, 110–117.

Wang, N., Arling, M., Hoerster, G., Ryan, L., Wu, E., Lowe, K., Gordon-Kamm, W., Jones, T.J., Chilcoat, N.D. and Anand, A. (2020) An Efficient Gene Excision System in Maize. Frontiers in Plant Science, 11.

